# Geographic distribution of terpenoid chemotypes in *Tanacetum vulgare* mediates tansy aphid occurrence and abundance

**DOI:** 10.1101/2023.06.19.545570

**Authors:** Humay Rahimova, Annika Neuhaus-Harr, Mary V. Clancy, Yuan Guo, Robert R. Junker, Lina Ojeda-Prieto, Hampus Petrén, Matthias Senft, Sharon E. Zytynska, Wolfgang W. Weisser, Robin Heinen, Jörg-Peter Schnitzler

**Affiliations:** Research Unit Environmental Simulation, Helmholtz Munich, D-85764 Neuherberg, Germany; Terrestrial Ecology Research Group, Department of Life Science Systems, School of Life Sciences, Technical University of Munich, D-85354 Freising, Germany; Fundamental and Applied Research in Chemical Ecology, Institute of Biology, University of Neuchâtel, 2000 Neuchâtel, Switzerland; Institute of Plant Protection, Beijing Academy of Agriculture and Forestry Sciences, 100097, Beijing, China; Evolutionary Ecology of Plants, Department of Biology, Philipps-University Marburg, 35043 Marburg, Germany; Julius Kühn Institute (JKI) – Federal Research Centre for Cultivated Plants, Data Processing Department, Stahnsdorfer Damm 81, 14532 Kleinmachnow, Germany; Department of Evolution, Ecology, and Behaviour, Institute of Infection, Veterinary and Ecological Sciences, University of Liverpool, L69 3BX Liverpool, United Kingdom

**Keywords:** chemotype, chemical diversity, geographical gradients, specialized metabolites, *Metopeurum fuscoviride Lasius niger*, *Formica rufa*, *Myrmica rubra*

## Abstract

**Aim:** Intraspecific variations of specialized metabolites in plants, such as terpenoids, are used to determine chemotypes. Tansy (*Tanacetum vulgare* L.) exhibits diverse terpenoid profiles, that affect insect communities. However, it is not fully known whether patterns of their chemical composition and associated insects vary on a large scale. Here, we investigated the geographic distribution of mono- and sesquiterpenoid chemotypes in tansy leaves and the effects of these chemotypes on colonization by insect communities across Germany.

**Methods:** We sampled tansy leaves from 26 sites along a north-south and west-east transect in Germany. Leaves from ten plants with and five plants without aphids was collected from each site. Hexane-extracted metabolites from leaf tissues were analysed by gas chromatography-mass spectrometry (GC-MS). Plant morphological traits, aphid occurrence and abundance, and occurrence of ants were recorded. The effect of plant chemotype, plant morphological parameters, and site parameters such as temperature and precipitation on insect occurrences were analysed.

**Results:** Plants clustered into four monoterpenoid and four sesquiterpenoid chemotype classes. Monoterpene classes differed in their latitudinal distribution, whereas sesquiterpenes were more evenly distributed across the transect. Aphid and ant occurrence were influenced by monoterpenoids and specific traits. Plants of monoterpenoid class 1 were colonized by *Metopeurum fuscoviride* and ants significantly more often than expected by chance compared to plants from monoterpenoid class 4. Aphid abundance was negatively affected by host plant height, and increasing average annual temperature positively influenced the occurrence of ants.

**Conclusion:** We found significant geographic differences in the chemodiversity of tansy and show that monoterpenoids affect aphid and ant occurrence, while host plant height can influence aphid abundance. We show that geographic variation in plant chemistry and morphology influences insect communities’ assemblage on tansy plants.

## 1.0 Introduction

Biogeography combines biological processes with the distribution of organisms (Tivy, 2018). Geographic location affects plant communities not only at interspecific levels but also at intraspecific levels (Moreira et al., 2012). One highly variable intraspecific plant trait can be the complex of specialized chemical compounds that vary considerably within plant families and even within the same plant species (Kessler & Kalske, 2018; Kleine & Müller, 2011). Individuals of the same plant species can be classified into different chemotypes characterized by their specialized metabolites. For instance, the abundance of pyrrolizidine alkaloids in *Senecio jacobaea* (Macel & Klinkhamer, 2010) and the abundance of specific monoterpenoids in *Pinus banksiana* (Taft et al., 2015), *Melaleuca alternifolia* (Bustos-Segura et al., 2017) and *Gossypium hirsutum* (Clancy et al., 2023) have been used for the classification of chemotypes. These primary and specialized metabolites in plants have many ecological functions and can strongly influence ecological interactions such as the attraction of herbivore predators, pollinators, and mycorrhizal fungi, defence against herbivores and pathogens, communication with other plants, and protection against UV-B radiation and drought (Dicke et al., 2009; Dixon & Paiva, 1995; Grof-Tisza et al., 2022; Mofikoya et al., 2019).

When biotic stress occurs, such as insect herbivory, various plant defence pathways can be activated, and a mixture of volatile compounds, including mono- and sesquiterpenoids, are induced and released (Niinemets et al., 2013). Moreover, volatiles serve as informational cues, e.g., for attracting natural enemies of herbivores (Baldwin, 2010; Heil & Bueno, 2007) and hence act indirectly as herbivore defence. However, plant morphology can also shape insect communities in addition to plant-specialized metabolites. For instance, in the perennial shrub *Baccharis pilularis*, plant architecture affected the composition of herbivore communities and morphology correlated with herbivory levels (Rudgers & Whitney, 2006). Furthermore, a plant’s surrounding vegetation can shape its volatile emissions and influence insect herbivores (Kigathi et al., 2019; Ziaja & Müller, 2023). Still, how specialized metabolites, morphological traits, the surrounding environment, and biogeographic differences, individually and combined, influence plant-insect interactions are not fully understood.

Tansy, *Tanacetum vulgare* L. (*Asteraceae*), is an aromatic herb endemic to Eurasia that exhibits a considerable variation in its terpenoid composition (Clancy et al., 2016; Keskitalo et al., 2001; Kleine & Müller, 2011). Terpenoids (isoprenoids) represent a large group of plant-specialized metabolites whose backbones consist of two common five-carbon isoprene units (Rosenkranz & Schnitzler, 2016). A widespread range of terpenoid synthases and subsequent modifying enzymes lead to numerous monoterpenoids and sesquiterpenoids (Degenhardt et al., 2009; Lange & Srividya, 2019). These terpenoids can be stored in glandular trichomes on the leaf surface in tansy (Guerreiro et al., 2016), or they can be induced and immediately emitted through biotic stresses such as herbivory (Clancy et al., 2016). The blends of the stored terpenoids vary between individuals, and tansy plants can be classified into different chemotypes according to those differences (chemical phenotypes; Kleine & Müller, 2011). For instance, tansy chemotypes characterized by the dominance of volatile terpenoids such as camphor, β-thujone, α-thujone, artemisia ketone, and borneol influenced the associated insect communities within a single field site (Kleine & Müller, 2011; Clancy et al., 2016). In addition to the volatile terpenoid pattern of tansy on a small scale, non-volatile metabolomic profiles can influence the abundance of aphids in the field (Clancy et al., 2018) and even shape the local genetic population structure of the herbivore (Zytynska et al., 2019). These discoveries highlight the role of intraspecific chemical diversity within tansy and its specialized aphids.

Aphids are sap-sucking plant-parasitic insects comprising up to 4000 species (Eastop, 1986). Most aphids have a single host-plant species (i.e., monophagous; Loxdale & Balog, 2018) and are exposed to many external forces that significantly impact their populations, such as predation and parasitism, environmental conditions, geography, and climate (Loxdale & Balog, 2018; Eastop, 1986). Furthermore, plant-related variables, such as host-plant chemistry, have strong effects on the behaviour and abundance of aphids. The plant chemical bouquet, for example, can determine aphid colonization (Neuhaus-Harr et al., 2023) or alter the predation rates on aphids (Linhart et al., 2005; Stadler, 2004). However, while there are abundant studies on large-scale variation in insect-plant interactions (Hortal et al., 2010; Rand & Louda, 2006; Tscharntke & Brandl, 2004), only a few have linked this variation to variation in plant chemistry (e.g., Berenbaum & Zangerl, 1998; Watt et al., 1997). This is despite the fact that chemical variation in a single host plant across space is common (Kessler & Kalske, 2018; Wetzel & Whitehead, 2020). As plant chemistry differs across geographical ranges, this may have ecological implications for associated insects.

In this study, we assess terpenoid variation in tansy on a regional scale and investigate consequences for plant-insect interactions. We carried out a thorough investigation of tansy along a north-south and west-east gradient in Germany, in which plant leaf samples, plant morphological traits, information on aphid abundance, ant species occurrence, abiotic site parameters, and the coordinates of the sampling sites were collected to answer the following questions:

1. How do terpenoid compounds in tansy cluster in chemotypes across a northwest-southeast transect in Germany? Moreover, are mono- and sesquiterpenoid chemotypes linked to one another?
2. How do chemotypes, plant architecture, and site variables affect aphid and ant occurrence and aphid abundance on plants?

We hypothesize that chemotypes of tansy derived from mono- or sesquiterpenoid contents are not associated with each other and that there are geographical differences in the distribution of this intraspecific variation of tansy in Germany. Moreover, we predict that chemotype and plant architecture significantly affect associated insect communities.

## 2.0 Material and Methods

### 2.1 Sampling of tansy plant populations and insect community

Tansy plants were sampled from 26 sites along a northwest-southeast transect in Germany (Fig. 1). The GPS coordinates of each sampling location were recorded on-site, with sites mainly located by country roads, train tracks, and agricultural edges. The sampling site around Bremen was the most northern site chosen (UTM Lon: 502110, Lat: 5875217), while the sampled site near Freising was the most southern one (UTM Lon: 718846, Lat: 5365982), spanning roughly 700 kilometres across Germany. The site near Bielefeld was the most western (UTM Lon: 476299, Lat: 5828202), while Leipzig was the most eastern sampling site (UTM Lon: 469284, Lat: 5692446), covering roughly 300 kilometres across Germany. The sampling survey took place from the 23^rd^ of June to the 23^rd^ of July in 2014. From each site, ten plants colonized by the specialized herbivore aphid *Metopeurum fuscoviride* (Stroyan) (minimum three stems occupied), and five aphid-free plants were selected for ecological analysis. For each plant, the presence of ants, as well as the ant species (*Formica rufa* L.*, Lasius niger* L., and *Myrmica rubra* L.) were recorded or marked as “unknown species” when they could not be identified. For plants with aphids, aphid abundance was calculated by counting the number of colonies, and estimating the size of each colony (XS: <10 aphids, S: 10-50 aphids, M: 50-200 aphids, L: >200 aphids). A subset of plants was randomly selected for chemical analysis.

**Figure 1:**
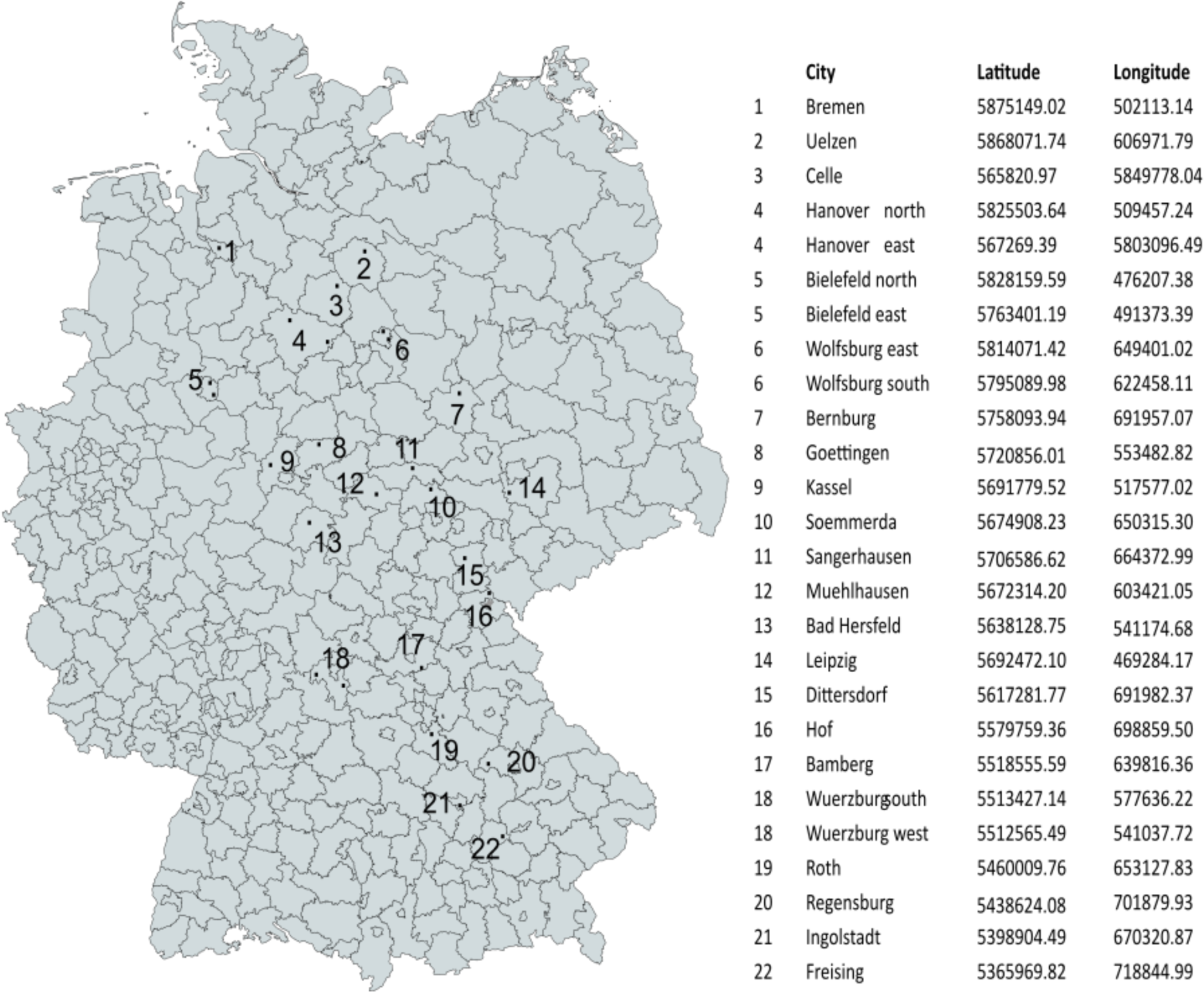
Map of the German cities near to the tansy sampling sites (26 sites were sampled across 22 districts) and their latitudinal and longitudinal values according to the UTM system.

### 2.2 Plant morphological measurements and abiotic site parameters

Plant morphological traits and geographic locations were recorded for each plant. Plant morphological traits included (1) the height of the tallest stem; (2) the number of stems; (3) the plant’s diameter at its widest width; (4) and the relative height of the tansy plant (higher/similar/lower than the surrounding vegetation). In addition, (5) the estimation of the percentage of grass, small herbs (<50 cm), tall herbs (>50 cm), shrubs, and bare ground within a two-meter radius were assessed, and as derived parameters, (6) the volume (radius^2^ * π * height), (7) the plant bushiness (plant volume divided by the number of stems), and (8) the emission potential of volatiles (volume * total terpenoid concentration) were estimated. Abiotic site parameters included the annual precipitation (reflecting the sum of rain over a year in mm) and the average annual temperature (in Celsius degrees) in every region. They were obtained from the German weather service (Kaspar, 2023).

### 2.3 Hexane extraction of terpenoids and GC-MS analysis

Leaf material collection and extraction were performed as described by Clancy et al. (2016). The detected compounds were identified by comparison of the mass spectra using the National Institute of Standards and Technology (NIST), Mass Spectral Library (NIST 11) and Wiley 275 GC/MS Library (Wiley, New York), and confirmed by comparison of the Kovats retention indices as reported by Guo et al., (2019, 2020) based on chromatography retention times of a saturated alkane mixture (C9 – C25; Sigma-Aldrich, Taufkirchen, Germany). The potential changes in the GC–MS sensitivity were corrected by normalizing to the internal standard (monoterpene δ-2-carene). The compounds were quantified using the external standards: sabinene, α-pinene, linalool, methylsalicylate, β-caryophyllene, α-humulene, geraniol, and bornyl acetate. The chemical structure of the identified compounds was sketched using ChemDraw professional (ChemDraw 21.0.0).

### 2.4 Statistical analysis

#### 2.4.1 Clustering plants into chemotypes

The plants were clustered into classes separately according to their monoterpenoid or sesquiterpenoid profiles by using the “*hclust*” function with the “*ward.D2*” method of correlation distance in the R “*factoextra*” package (Kassambara & Mundt, 2020a). The “statistical meta-analysis” function in the online software MetaboAnalyst 5.0 (Pang et al., 2021) was used to compute the heatmap of the monoterpenoid and sesquiterpenoid compounds contributing to the differentiation of the classes. Discriminant analyses of principal components (DAPC) were applied to infer the separation of monoterpenoid and sesquiterpenoid chemotype classes. The number of retained principal components (PCs) was determined by cross-validation using the “*xvalDapc*” function in the “*adegenet*” package v2.1.1 (Jombart, 2008) in R.

Additionally, to compare the relationship between mono- and sesquiterpenoid classes, a tanglegram of both dendrogram trees was obtained by using the *“dendlist”* function in the *“dendextend”* package (Galili, 2015). We also tested the associations between the selected monoterpenoids using correlation analysis with a Spearman method at a 99% confidence interval by using in *“ggpubr”* package (Kassambara, 2020b).

All other graphs were made using the package ggplot2 in R (Wickham, 2016). For better resolution, the resulting images were edited using the image processing software “*Inkscape*” ® (version 1.1.1).

#### 2.4.2 Phytochemical analysis

To compare the phytochemical diversity for the chemotype classes, we calculated the functional Hill diversity (FHD) for all samples, separately for monoterpenoids and sesquiterpenoids, using the “*chemodiv*” R package (Petrén et al., 2023a). The functional Hill diversity (FHD) was calculated at diversity orders from q = 0 to q = 3. For increasing q-values, the measure puts more weight on abundant compounds; at q = 0, the relative abundances of compounds are not taken into account; at q = 1, the weight is proportional to their abundance, and at q > 1, more weight is put on abundant compounds of which the upper limit is set to q = 3. Dissimilarities between compounds were calculated based on PubChem Fingerprints (Kim et al., 2021), which quantify dissimilarities based on the structural properties of the molecules. Each compound’s chemical identifiers (SMILES and InChIKey) were extracted from the PubChem open database (https://pubchem.ncbi.nlm.nih.gov).

#### 2.4.3 Distribution of terpenoid chemotypes across a geographical transect in Germany

We carried out a permutational multivariate analysis of variance (PERMANOVA, “*adonis2*” function in “*vegan*” R package; Oksanen et al., 2020) using chemical distance matrices (999 permutations, Bray-Curtis method) versus the latitude and longitude across each pairwise combination.

#### 2.4.4 Morphological differences across chemotypes and a geographical transect

To test whether plant morphology differed across the latitudinal or longitudinal gradient in Germany and whether it differed between chemotype classes, we used a one-factorial ANOVA. We tested the number of stems, plant volume, emission potential, plant height, radius, and bushiness.

#### 2.4.5 Effect of chemotypes on associated insect community

To test whether aphid occurrence or ant occurrence was influenced by chemotypes, plant morphology, and site variables, we set up a generalized linear model (GLMs) with binomial distribution when the response variable was occupancy (1/0). For aphid abundance, the number of colonies per plant was multiplied with the minimum number of aphids in each colony category. The response variable, the number of aphids, was log-transformed to ensure that the assumption of normality was met, and a linear model with a normal distribution (LM) was used. The “*emmeans”* R package with Tukey adjustment was used to assess post-hoc pair-wise comparisons among factor levels following model fit (Russell, 2021).

Prior to running the generalized models and to rule out multicollinearity, we excluded all variables with a variance inflation factor higher than five (“*VIF*” function in “*car*” R package, Fox & Weisberg, 2019). We omitted “plant radius”, “plant volume”, and “total terpenoid concentration”. As predictor variables, we included “monoterpenoid class”, “sesquiterpenoid class”, “monoterpenoid concentration”, “sesquiterpenoid concentration”, “emission potential”, “bushiness”, “height of surrounding vegetation”, “plant height”, “the number of stems”, “annual temperature at the site”, “annual precipitation at the sampled site”, “latitude” and “longitude” in all models.

To test whether the FHD affected aphid occurrence and abundance, we used a one-way analysis of variance test. Functional Hill diversity of mono- and sesquiterpenoids were log-transformed to meet normality assumptions. Statistical models were carried out using R (version 4.1.3; R Core Team 2022).

## 3. Results

### 3.1 Monoterpenoid and sesquiterpenoid tansy chemotypes

#### 3.1.1 Tansy monoterpenoid and sesquiterpenoid chemotypes

We analysed 278 plants and identified 30 monoterpenoids and 21 sesquiterpenoids. The molecular schemes of some mono- and sesquiterpenoids are depicted in Fig. 2a and 3a. Plants clustered into four distinct monoterpenoid (Fig. 2b) and four sesquiterpenoid chemotype classes (Fig. 3b). The eastern Hanover sampling site was excluded for further analyses because two unknown sesquiterpenoid compounds found there could not be annotated by library search and Kovats index comparisons, and this site specifically showed exceedingly high concentrations of β-thujone and bicyclosesquiphellandrene, causing severe outliers (Fig. S1).

**Figure 2:**
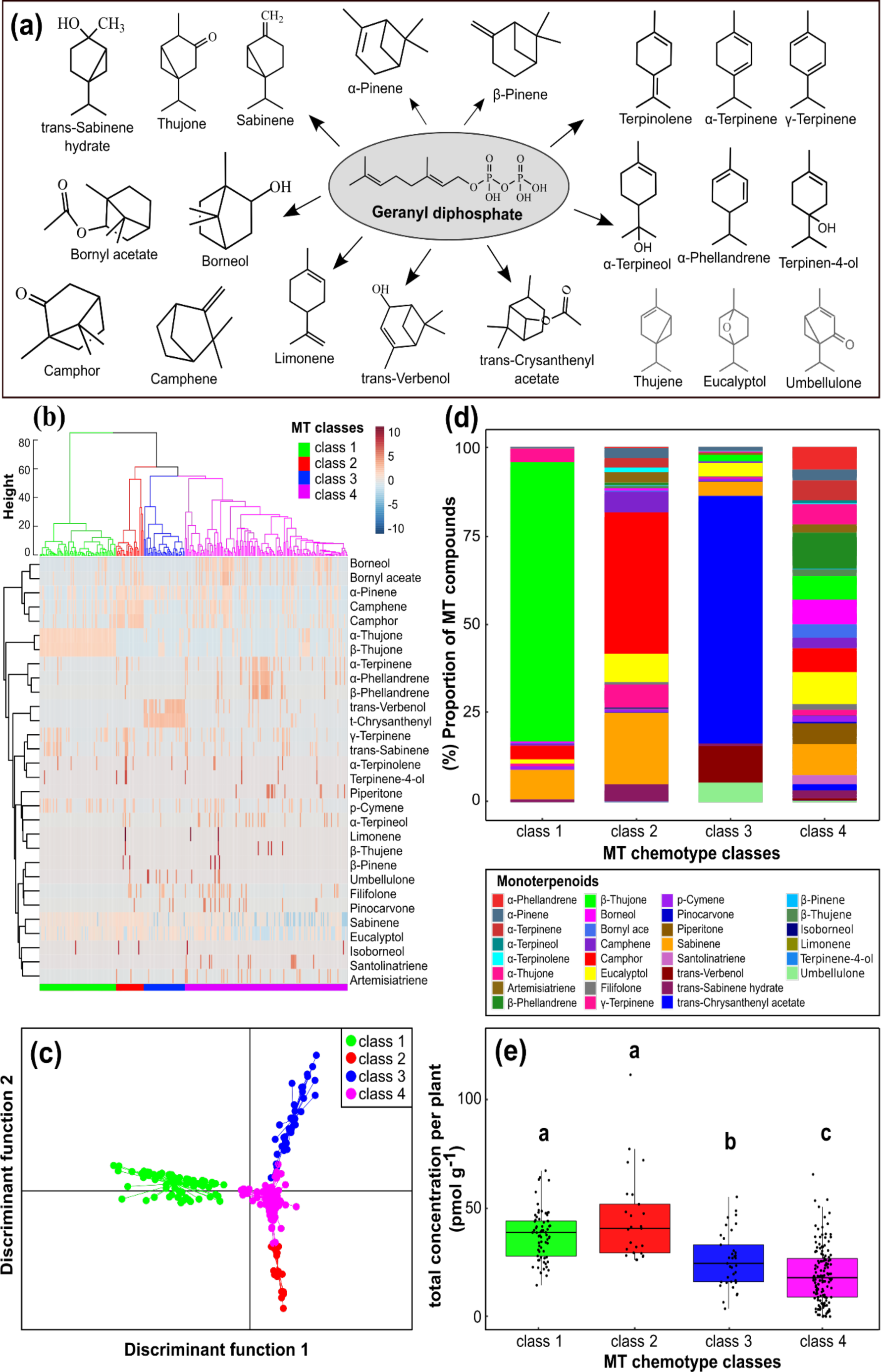
Visualization of the monoterpenoid (MT) chemotype classes using different statistical approaches. (a) Schematic illustration of Monoterpenoid products synthesized from Geranyl diphosphate via carbocationic reactions mechanism. The compounds that are biosynthetically linked stand in the same row, e.g. sabinene, thujone and trans-sabinene hydrate (b) Hierarchical Cluster Analysis of monoterpenoid compounds across 278 individual plants. Four main classes were identified and each cluster is highlighted by a different color; class 1 green, class 2 red, class 3 blue, and class 4 magenta. The variety and separating features of monoterpenoids found in each class is displayed in the heatmap. (c) Discriminant analysis of principal components plot shows separations among the MT chemotype classes. (d) Real examples as well as the proportional (in percentage) composition of each chemotype are provided in a stacked barplot. Data is normalized to logarithmic scale and Pareto matrix. (e) Total concentration of monoterpenoids with significant differences (Tukey test, p < 0.05) indicated by letters.

**Figure 3:**
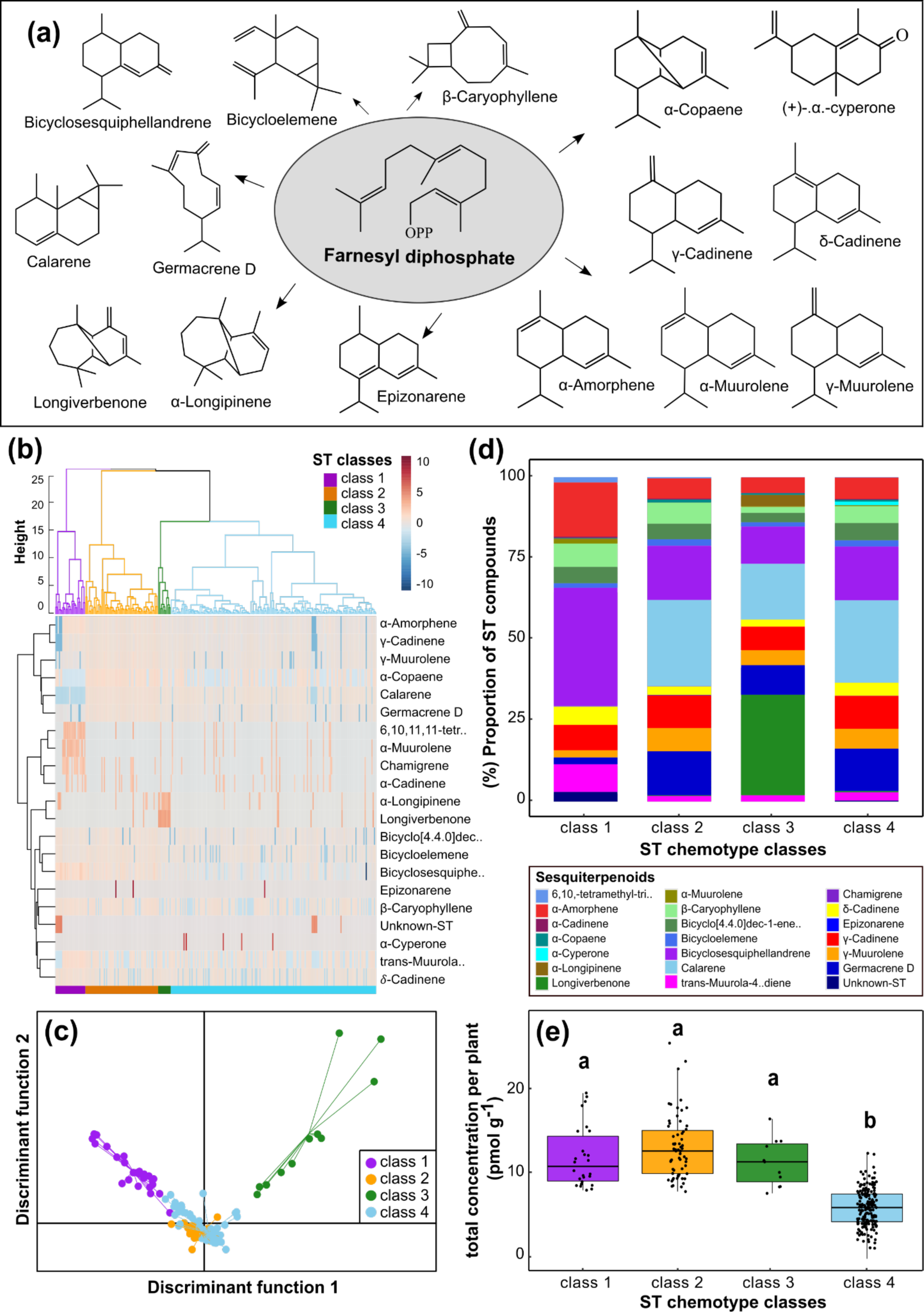
(a) Schematic illustration of sesquiterpenoid (ST) products synthesized from farnesyl diphosphate. All the sesquiterpenoids are the main products of farnesyl diphosphate (b) Hierarchical Cluster Analysis of sesquiterpenoid compounds across 278 plants; Four main classes were classified – class 1 purple, class 2 orange, class 3 forestgreen, and class 4 lightblue. The variation of the sesquiterpenoid compounds of each class is depicted in the heatmap. Data is logarithmically transferred and Pareto scaled. (c) Discriminant analysis of principal components indicates separations among the sesquiterpenoid classes. (d) Proportional composition of the compounds, provided in percentage for each class. (e) Total concentration of sesquiterpenoids with significant differences (Tukey test, p < 0.05) indicated by the letters on the top.

A heatmap of monoterpenoids shows the separating features of α- and β-thujone in class 1, camphor and camphene in class 2, trans-verbenol and trans-chrysanthenyl acetate in class 3, and α-terpinene, α- and β-phellandrene in class 4 (Fig. 2b). Monoterpenoid classes comprised 69, 25, 37, and 147 plants, for classes 1, 2, 3, and 4, respectively. Discriminant analysis of principal components models (DAPC) showed a discrimination among the different monoterpenoid classes (Fig. 2c). Specifically, class 1 was dominated by β-thujone (approximately 75%), class 2 was defined by a mixture of approximately 40% camphor and 20% sabinene, class 3 was dominated by trans-chrysanthenyl acetate (approximately 60%), and class 4 comprised a group of plants with a mix of compounds (Fig. 2d). The total concentration of monoterpenoids varied from 0.02 to 112 pmol g^-1^ leaf fresh with classes 1 and 2 having the highest monoterpenoid concentrations and class 4 having the lowest (Fig. 2e). Additionally, we found that some monoterpenoids were closely linked to each other. Regardless of the chemotype categorization, camphene and camphor concentrations were associated in monoterpenoid classes 1, 2, and 4 (R^2^ = 0.42, p < 0.001; Fig. S2a), and borneol and bornyl acetate concentrations showed a positive correlation (R^2^ = 0.50, p < 0.001; Fig. S2b) among the many plants.

Plants clustered into four sesquiterpenoid chemotype classes, with the compound pattern of each sesquiterpenoid class presented in a heatmap (Fig. 3b). Here, 26, 63, 11, and 175 plants were classified into classes 1, 2, 3, and 4, respectively. A DAPC model showed a discrimination among the sesquiterpenoid classes (Fig. 3c). In contrast to distinct monoterpenoid composition between classes, sesquiterpenoid classes showed a general presence of bicyclosesquiphellandrene, β-caryophyllene, γ-muurolene, and α-amorphene (Fig. 3d). The plants belonging to sesquiterpenoid class 1 were characterized by the highest proportion of bicyclosesquiphellandrene (36%). In comparison, calarene was almost absent (0.05%). Classes 2 and 4 showed a profile of calarene (25%), germacrene D (13%), and γ-cadinene (10%). Furthermore, although sesquiterpenoid class 3 contained fewer individuals, it was predominantly formed by longiverbenone (31%). The total concentration of sesquiterpenoids ranged from 1.3 to 25.5 pmol g^-1^ (Fig. 3e).

The tanglegram of monoterpenoid and sesquiterpenoid chemotypes across all tansy individuals showed no clear link between monoterpenoid and sesquiterpenoid profiles (Fig. 4a). Some monoterpenoid individuals were tangled with different sesquiterpenoid classes. For instance, most plants belonging to monoterpenoid class 4 were linked to sesquiterpenoid classes 1 and 2, while monoterpenoid classes 1 and 3 were mainly found related to sesquiterpenoid class 4.

**Figure 4:**
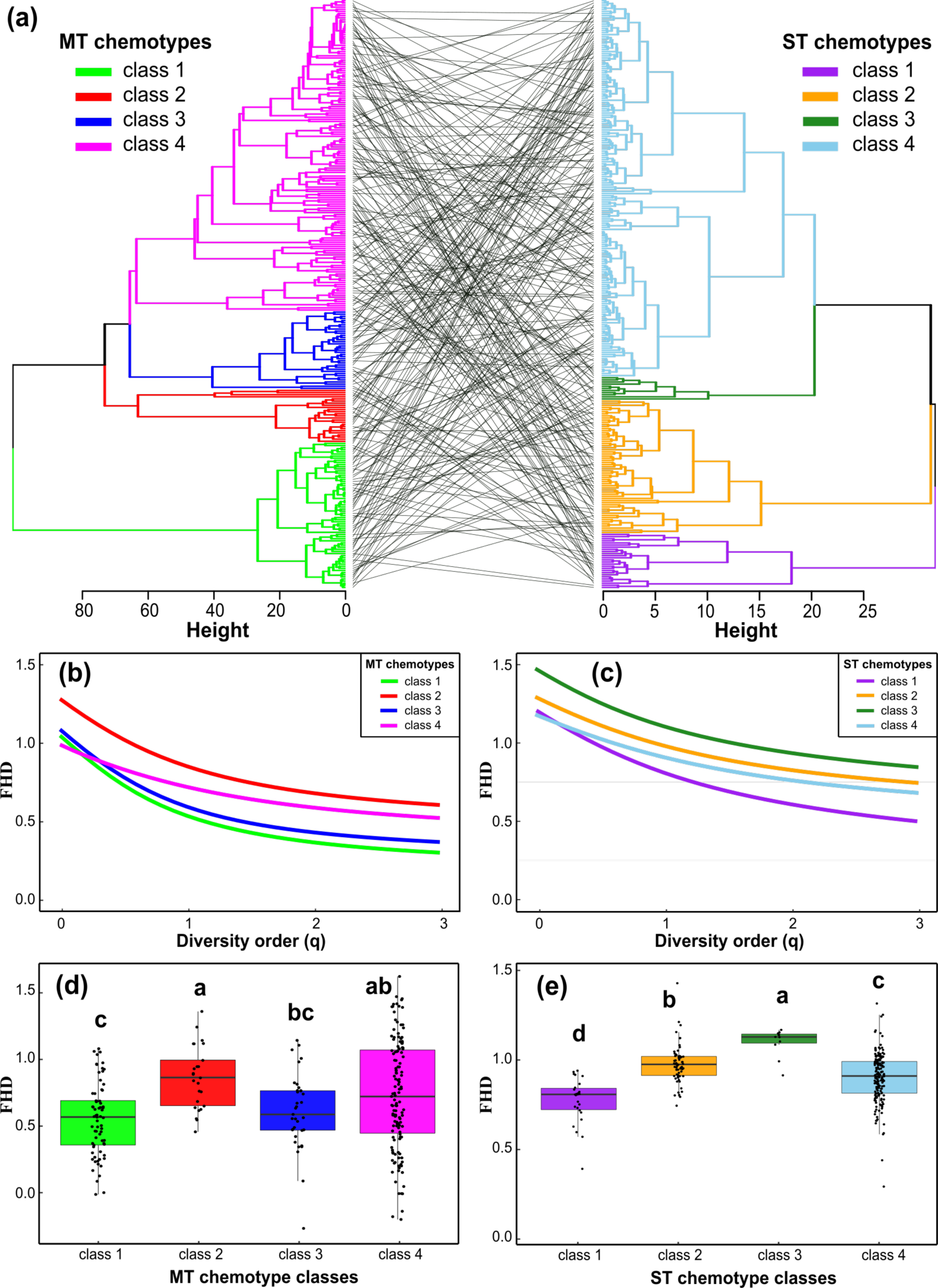
(a) A tanglegram of the chemotype trees demonstrating that monoterpenoid (MT; on the left) and sesquiterpenoid (ST; on the right) chemotype classes are not linked across the 278 individuals. The diversity profile shows the functional Hill diversity (FHD) at diversity orders from q = 0 to q = 3 for MT (b) and ST (c) chemotype classes. For increasing q-values, the measure is less sensitive to the relative abundances of compounds; at q = 0 the relative abundances of compounds are not taken into account; at q = 1 equal weight is put on all compounds. The boxplots show variation in FHD for the MT (d) and ST (e) classes in detail for q = 1. Note that values are log-transformed in figures and analyses. Significant differences (p < 0.05) between classes are indicated by the letters above the boxes. Mean number of monoterpenoids per sample: class 1 = 5.9, class 2 = 7.2, class 3 = 5.7, class 4 = 5.6. Mean number of sesquiterpenoids per sample: class 1 = 10.3, class 2 = 10.7, class 3 = 12.0, class 4 = 9.7.

#### 3.1.2 Phytochemical diversity metrics across chemotype classes

Quantifying the functional Hill diversity enabled us to study the diversity of tansy monoterpenoid and sesquiterpenoid chemotype classes in novel ways that differ in the weight placed on abundant compounds through compound richness, evenness and disparity (Petrén et al., 2023b). Functional Hill diversity among monoterpenoid classes was the lowest for classes 1 and 3, which was lowest at higher orders of q, emphasizing the role of abundant compounds in these classes. Monoterpenoid class 2 had the highest functional Hill diversity, but a rather deep curve, again indicating that abundant compounds are likely important in this class. Class 4 had an intermediate mean functional Hill diversity, with a shallow curve (Fig. 4b, d), which indicates that dominant compounds play a less important role in this class, which is indicated by a high evenness in the diversity profile (Fig. 2d). Notably, the diversity at the level of chemotypes, rather than individual plants, appeared to be higher in class 4, as an effect of larger differences between samples in their composition.

Within the sesquiterpenoid classes, class 1 and 3 had a deeper curve (Fig. 4c), with low and high functional Hill diversity (Fig. 4e), emphasizing a role of dominant compounds in shaping the functional Hill diversity (Fig 3d). Sesquiterpenoid classes 2 and 4 had more similar and shallower curves with intermediate mean functional Hill diversity (Fig. 4c, e), indicating a more even sesquiterpenoid profile (Fig. 3d). Overall, the functional Hill diversity of sesquiterpenoids differed significantly between all classes (Fig. 4e).

#### 3.1.3 North-south and west-east gradient in monoterpenoid and sesquiterpenoid chemotypes

The distribution of tansy monoterpenoid classes differed significantly across Germany. Monoterpenoid classes 1 and 2 were found more frequently in the east and south, while monoterpenoid classes 3 and 4 were more frequently observed in North- and West-Germany (Fig. 5a). A PERMANOVA test showed that monoterpenoid compositions significantly varied depending on the latitude (R^2^ = 0.01; p < 0.002; Fig. 5c) and longitude (R^2^ = 0.01; p < 0.001; Fig. 5e, Table S1) coordinates, even though the explained variance (R^2^) is low. Biosynthetically linked monoterpenoids, such as β-thujone and sabinene, camphor, and camphene, increased substantially with decreasing latitude and increasing longitude towards the far south. Contrastingly, trans-crysanthenyl acetate and trans-verbenol showed an opposite trend, with the highest concentration reported in plants at high latitudes towards more northern sites (Fig. S3).

**Figure 5:**
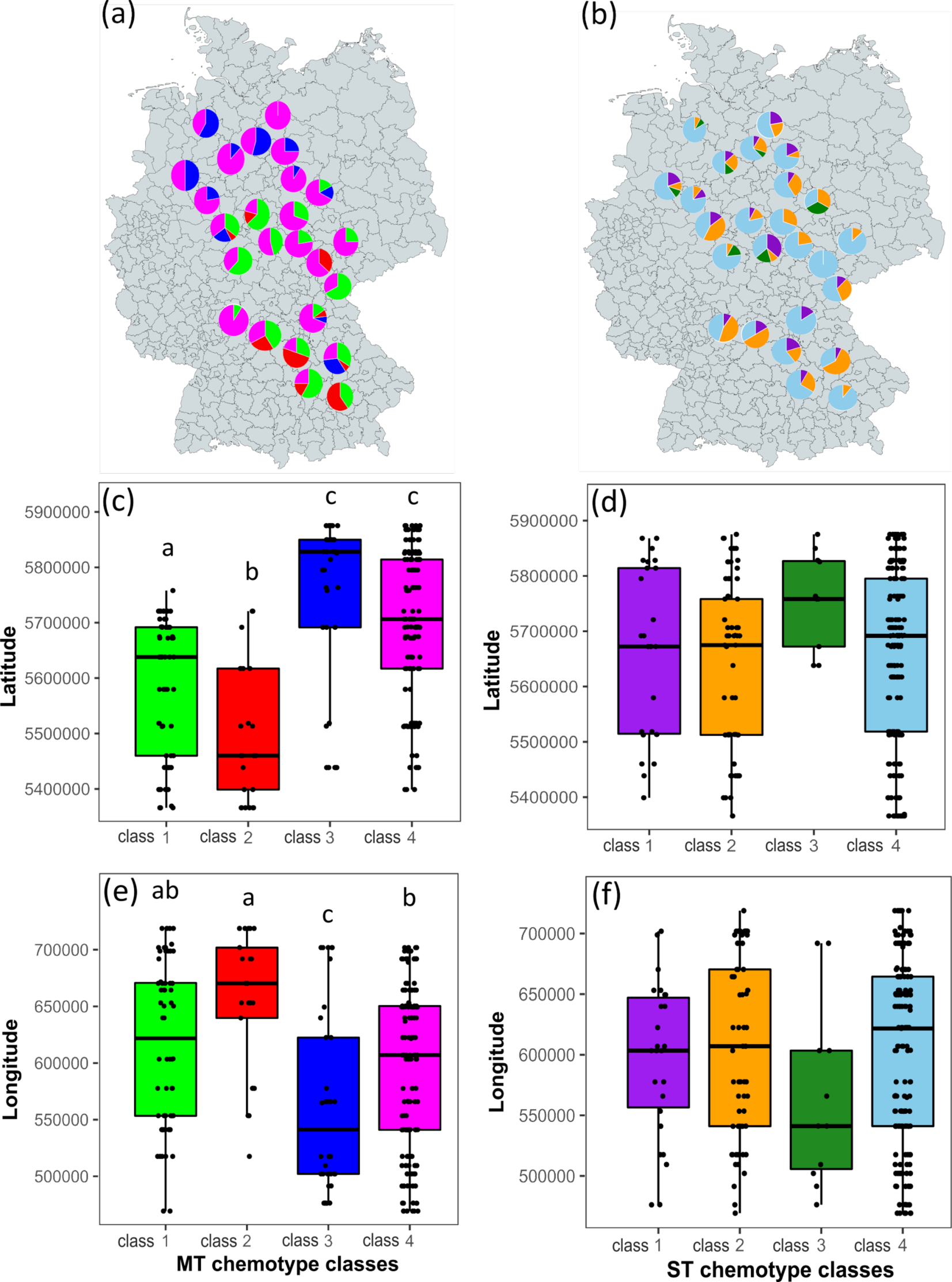
Proportion of monoterpenoid (MT) and sesquiterpenoid (ST) classes within each sampled site. Monoterpenoid classes are color-coded as following: green - class 1, red – class 2, blue – class 3, magenta – class 4 (a). Sesquiterpenoid classes are color-coded as following: purple – class 1, orange – class 2, dark green – class 3, light blue – class 4 (b); monterpenoid classes found over different latitude (c) and longitude (e) and sesquiterpenoid classes found over different latitude (d) and longitude (f); significant differences are indicated on top of the boxplots (p < 0.05).

In contrast, sesquiterpenoid classes were more homogeneously distributed across Germany (Fig 5b), indicating independence of the geographic effects on mono- and sesquiterpenoid chemotypes (Fig 5d, f). Sesquiterpenoid compositions did not significantly differ in their geographic distribution (lat.: R^2^ = 0.002; p = 0.56; lon.: R^2^ = 0.002; p = 0.58; Table S1).

### 3.2 Plant morphology differences between chemotypes and across Germany

The number of stems per plant differed marginally significantly between monoterpenoid classes (F_(3)_ = 2.55, p = 0.056; Table S2). A post-hoc test showed that the number of stems was lower in plants from monoterpenoid class 1 compared to class 4 (Fig 6a). Plant volume and emission potential differed significantly between plants of different monoterpenoid chemotypes (F_(3)_ = 2.71, p = 0.045; F_(3)_ = 8.71, p < 0.001; Table S2). Plant volume was significantly lower in plants belonging to monoterpenoid class 1 than in class 2, but neither differed from monoterpenoid classes 3 and 4 (Fig. 6b). Plants belonging to monoterpenoid class 2 had significantly higher emission potential than all other monoterpenoid classes (Fig. 6c).

**Figure 6:**
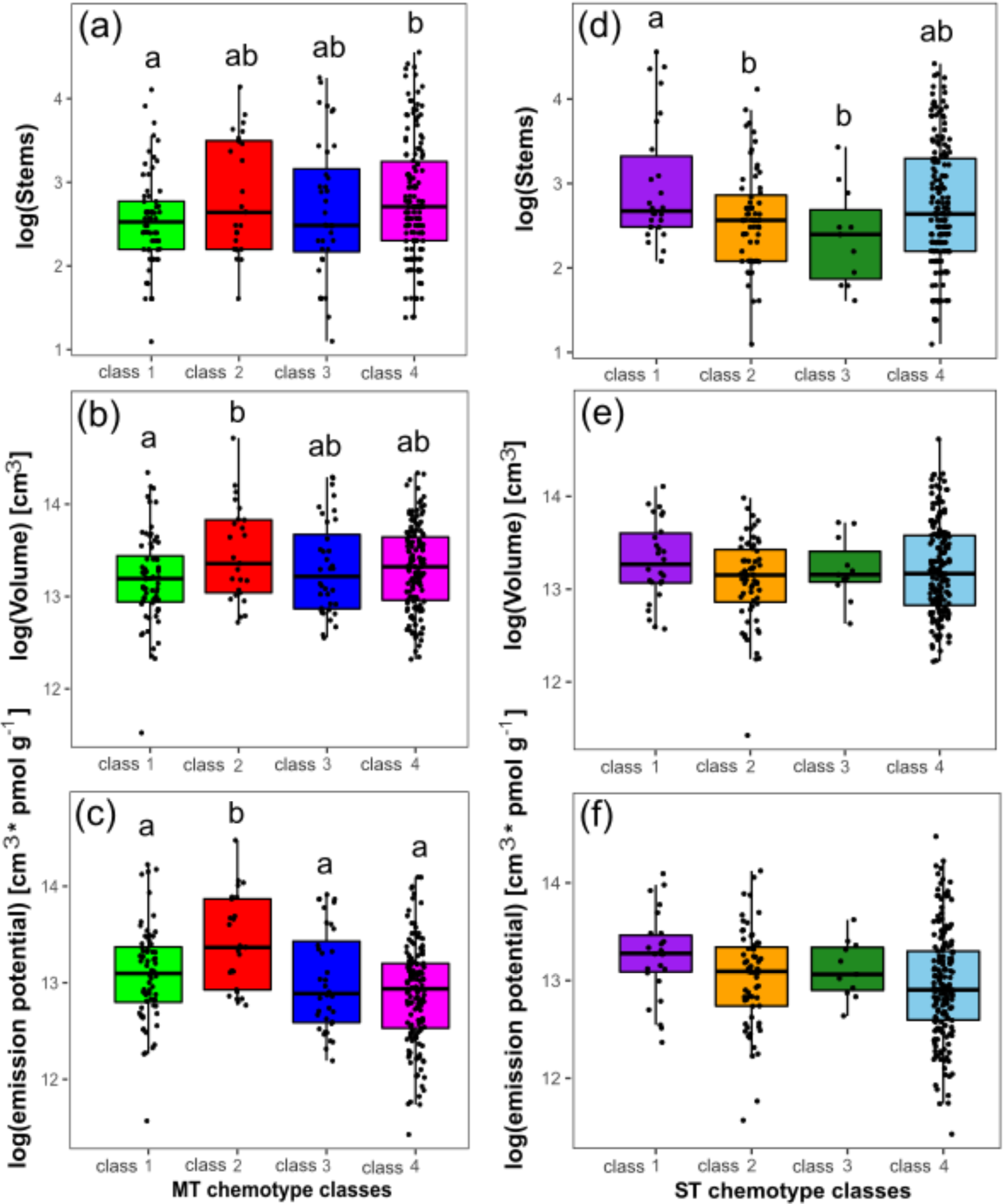
Plant traits with a significant difference across monoterpenoid classes: Number of stems (a), plant volume (b) and emission potential (c). Plant traits across different sesquiterpenoid classes: Number of stems (d), plant volume (e) and emission potential (f). Emission potential was calculated by the plant volume * the concentration of all terpenoid compounds in that specific plant. Degrees of freedom (DF), statistic (F-value) and p-value for plant trait variation across monoterpenoid and sesquiterpenoid classes using a one-factorial ANOVA are indicated in Table S3.

Sesquiterpenoid classes showed substantial differences in the number of stems (F_(3)_ = 3.69, p = 0.012; Table S2). Specifically, plants from sesquiterpenoid class 1 had a significantly higher number of stems than sesquiterpenoid class 2 and 3 (Fig. 6d). However, there were no differences in their volume or emission potential (Fig. 6e, f). Plant height, plant radius, and plant bushiness did not differ significantly across monoterpenoid nor sesquiterpenoid classes. F-statistics and p-values for all measured plant traits are in the appendix (Table S2).

Plant traits also varied across the geographical gradient. Plant height differed significantly along the latitudinal gradient (F_(3)_ = 10.20, p = 0.002, Table S3), with plants typically growing taller in the north. Plant bushiness differed across the longitudinal gradient (F_(3)_ = 7.35, p < 0.01, Table S3), with plants growing bushier in the west. Most plant variables, such as radius and number of stems, were positively correlated (i.e., higher plants tended to be bushier).

### 3.3 Effects of site conditions and plant chemical and morphological traits on insect tansy-associated community

Modelling the effects on aphid occurrence with a binomial GLM, we found that aphid occurrence was affected by chemical and morphological plant traits. Specifically, the monoterpene class (X^2^_(3)_ = 11.99, p = 0.007, Fig. 7a) and, marginally significantly, the number of stems (X^2^_(1)_ = 3.76, p = 0.053) influenced the occurrence of *M. fuscoviride* aphids (Table S4).

**Figure 7:**
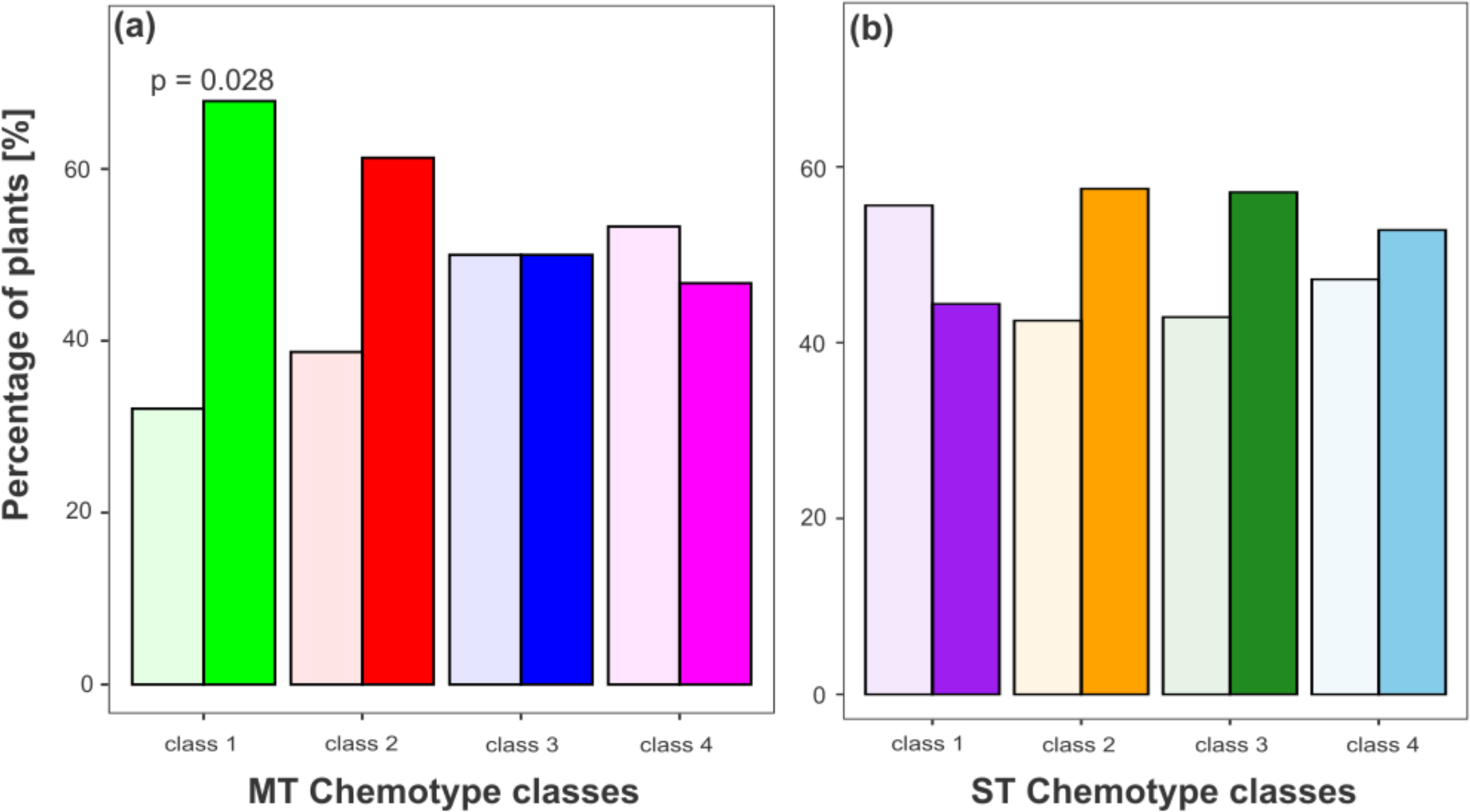
Bar charts indicate the percentage of plants without *M. fuscoviride* aphids (transparent) and with aphids (opaque) within monoterpenoid (MT) classes (a) and sesquiterpenoid (ST) classes (b). A binomial test showed that plants of monoterpenoid class 1 (a) were colonized by this aphid species significantly more often than expected by chance (binomial test: 95% conf. interval = 0.52 - 0.82, p = 0.028; **Table S4**). Sesquiterpenoid classes (b) did not influence aphid occurrence (**Table S1**, **Table S4**).

Sesquiterpenoid classes did not influence aphid occurrence (Fig. 7b, Table S4, S5). Opposed to this, a linear model with aphid abundance revealed that aphid abundance was not affected by monoterpenoid classes, but was marginally influenced by the height of the tansy plant (F_(1)_ = 3.17, p = 0.077; Table S4, S5), with *M. fuscoviride* aphids being less abundant on taller plants. However, when checking for correlations, we found that the abundance of *M. fuscoviride* was not correlated with any plant variables nor to single (dominant) compounds of the monoterpenoid classes (spearman correlation with 0.95-confidence level; Fig. S4). Even though the aphid occurrence responds to monoterpenoid classes, the functional hill diversity of neither the monoterpenoids nor the sesquiterpenoids affected *M. fuscoviride* presence or numbers significantly (Table S6).

Three species of ants were observed regularly in all sites and often on the same plant (*Formica rufa* L.*, Lasius niger* L., or *Myrmica rubra* L., Fig. S5a). A binomial GLM with ant presence as dependent variable showed that monoterpenoid class, but not sesquiterpenoid class, significantly affected probability of ant presence (X^2^_(3)_= 10.62, p = 0.014; Table S4, Fig S6a, b), and posthoc Tukey tests indicated that monoterpenoid class 1 had significantly higher probability of ant presence than monoterpenoid class 4. Furthermore, ants were more likely to be present at sites with higher temperatures (X^2^_(1)_ = 3.91, p = 0.048; Table S4, Fig. S7). This was independent of the species.

Parasitized aphids were commonly observed in all sites (Fig. S5b). However, we did not observe clear drivers of the probability of parasitisation except latitude and longitude. Specifically, parasitisation was more common in the southwest than in the northeastern sites (Fig. S5b).

## Discussion

We demonstrated that *Tanacetum vulgare* plants exhibit variation in distinct mono- and sesquiterpenoid chemotypes across a wide geographical range in Germany. Our results show that the chemical composition of monoterpenoids was significantly different across geographical coordinates, demonstrating that the monoterpenoids profile of tansy was more dissimilar with increasing geographical distance. While monoterpenoid chemotypes displayed different local dominance patterns, sesquiterpenoid chemotypes were homogeneously distributed across Germany. We further demonstrate that monoterpenoid classes, but not sesquiterpenoid classes, are involved in shaping aphid (*M. fuscoviride*) and ant *(L. niger*, *F. rufa*, and *M. rubra*) occurrence patterns. Furthermore, plant height affected the number of aphids while mean annual temperature had a positive influence on ant occurrence, suggesting that chemical, morphological and geographic factors structure the wider ecological community.

Tansy chemodiversity has been investigated in different geographical regions of Europe. For instance, a study from Finland revealed that Finland’s central and southern regions were home to tansy chemotypes with higher concentrations of camphor (Keskitalo, Pehu & Simon, 2001). Interestingly, we also found that plants from monoterpenoid class 2, which is dominated by camphor, were more frequent in southern Germany. Tansy seems to differ in its terpenoid profile between and within countries. For example, tansy plants from Finland showed a unique davadone D chemotype, while myrcene-tricyclene chemotypes were more common in the south and southwest compared to the rest of the country (Keskitalo, Pehu & Simon, 2001). Moreover, a study from Lithuania found that tansy exhibited different dominant compounds (such as eucalyptol, trans-thujone, and myrtenol) between different locations (Judzentiene & Mockute, 2005). In line with these findings, we observed plants from the β-thujone chemotype more prevalent and plants from the trans-chrysanthenyl acetate chemotype less prevalent in the south of Germany compared to the northern German cities. These findings suggest that differences in terpenoid profiles are common and likely increase with larger scales and that different regions bolster different dominance patterns of terpenoid compounds.

Our study found that on a German-wide scale, tansy plants could be grouped into distinct chemotypes using their mono- and sesquiterpenoid profiles. In this body of work, hierarchical cluster analysis revealed four monoterpenoid and four sesquiterpenoid classes that were not strongly associated with one another. This lack of alignment between individuals of mono- and sesquiterpenoid chemotypes strongly suggests differing biosynthesis pathways for these two compound classes. Sesquiterpenoids are generally produced through the cytosolic mevalonate pathway (MVA), whereas the plastidial methylerythritol phosphate (MEP) pathway yields multiple monoterpenoid products (Davis & Croteau, 2000). However, the MVA and MEP can provide isopentenyl diphosphate precursors for monoterpenoid and sesquiterpenoid biosynthesis (Dudareva et al., 2005). This may explain why some individuals of monoterpenoid chemotype classes link with sesquiterpenoid chemotype classes. The diversity of terpenoid compounds in plants is generated by terpene synthases, a diverse family of enzymes that catalyze terpenoid compounds from single substrates (Bohlmann et al., 1998). For instance borneol and bornyl acetate showed a significant positive correlation in the same plant where they both were active, indicating that they are likely produced from the synthase of the bornyl diphosphate enzyme (Fig. S2). A similar result was observed between camphor and camphene. Interestingly, the chemotypes also differed in their phytochemical diversity in compound richness, evenness, and dissimilarity, which may impact interactions between plants and insects (Whitehead et al. 2021; Neuhaus-Harr et al. 2023).

We assessed the impacts of monoterpenoid and sesquiterpenoid composition on interactions with a specialized insect herbivore, i.e., *Metopeurum fuscoviride*, and three species of ants (*Formica rufa* L., *Lasius niger* L., and *Myrmica rubra* L.). Plants belonging to monoterpenoid class 1 were significantly more likely to be colonized by aphids, whereas equal occupancy of plants was observed for all sesquiterpenoid classes. This finding is in line with other studies, since our monoterpenoid class 1 contains β-thujone as a dominant compound, which has been associated with an increased abundance of another tansy specialist aphid, *Macrosiphoniella tanacetaria* (Kleine & Müller, 2011). Interestingly, previous studies also found higher abundance and earlier colonization rates of *M. fuscoviride* on plants with camphor as dominant compound, which would resemble monoterpenoid class 2 in our study (Clancy et al., 2016; Senft et al., 2019). This could explain why we observed the tendency of higher aphid presence on monoterpenoid class 2 plants, even though this finding was not significant. However, other studies have shown that not only dominant but also minor compounds within a blend significantly affect plant-insect interactions (McCormick et al., 2014; Clancy et al., 2016).

Preference, and therefore presence, appears to be affected by terpenoids, a finding in line with the idea that volatile terpenoids serve as cues for finding host plants (Bruce, Wadhams & Woodcock, 2005; Ninkovic, Markovic & Rensing, 2021). It is possible that monoterpenoids are more helpful to aphids as cues for host plant identification than sesquiterpenoids, as plants exhibited much higher concentrations of monoterpenoids compared to sesquiterpenoids. Clancy et al. (2016) observed that the emission of terpenoids, presumably evaporated/released from glandular cells (Devrnja et al., 2021), affected *M. fuscoviride* colonization. Given the higher volatility of monoterpenoids (Mofikoya et al., 2019), higher concentrations could be expected in the near ambient air of the plants’ canopy. This fits with the observation that the influence of terpenoids on aphid presence on individual chemotypes is mainly determined by monoterpenoids and not by sesquiterpenoids. If monoterpenoids are used a host-finding cue, the significant low concentration of monoterpenoids in monoterpene class 4 could perhaps be the reason why we see a tendency of low aphid presence in these plants. Another reason, why mono- and not sesquiterpenoids could be used as host finding cues by aphids, could be the differentiation of the monoterpenoid profiles in terms of their (dominant) compounds. Profiles were very distinct in monoterpenoids, while the sesquiterpenoid classes were chemically more similar. Functional Hill diversity had no effect on aphid presence and abundance. Functionally related terpenoids seem to have the same effect on aphids compared to functionally unrelated terpenoids.

Furthermore, not only aphids, but also ants might use monoterpenoids as cues. As *M. fuscoviride* is a facultative ant-tended species that benefits strongly from mutualism (Flatt & Weisser, 2000), chemotypes might structure aphid colonization and population indirectly via ant preference. Hence, it is unsurprising that we found a higher ant occupancy of plants belonging to monoterpenoid class 1 compared to plants from monoterpenoid class 4, similar as in the aphids. Even though we could not confirm whether ant presence was shaping aphid presence, or vice versa, this suggests that plant chemotypes mediate the strong relationship between those insects in tansy (Mehrparvar et al., 2017). Even more so, it has been found that the presence of ants before aphid appearance led to a stronger likelihood of aphid colonization (Senft et al., 2018). This suggests that aphids may have an increased preference for plants of specific chemotypes (Neuhaus-Harr et al., 2023), but also that chemotypes could affect aphid survival and population growth, e.g., via interactions with ants (Mehrparvar et al., 2017). Furthermore, we recognized that geographically changing environmental factors affected the abundance of ants, as their abundance increased with the average annual temperature of the site. This has already been shown for Mediterranean ant species (Cerdá et al., 1998). For example*, Lasius niger*, known for its thermal tolerance and preference for temperatures in the 18 - 26 °C range, increases their foraging activity at higher temperatures (Blanchard et al., 2021), which could ultimately influence aphid presence and abundance. Hence, chemical cues, such as mono- or sesquiterpenoids might only be one factor shaping aphid communities.

Although we show an effect of chemotypes on plant occupancy by aphids, we did not find such links with aphid abundance. Furthermore, we did not observe links between individual terpenoid compounds and aphid abundance. Several studies now show links between terpenoid composition on aphid preference, and presence on a plant for various aphid species in this model system (Neuhaus-Harr et al. 2023). However, aphid presence on a plant can vary strongly in their abundance, from very few aphids to thousands. These differences in aphid abundance partially may occur after a plant is colonized, and hence can also be shaped by other factors than plant chemotype (Neuhaus-Harr et al. 2023), such as predation pressure (Martínez, Soler & Dicke, 2013; Senft et al., 2019) or resource availability. Indeed, it has been found that *coccinellids*, which predate heavily on aphids, are more abundant on plants with high β-thujone contents (Kleine & Müller, 2011), which might explain why aphids on plants with high β-thujone did not show higher abundances.

We also found that aphid abundance was affected by host plant height. Other morphological traits, such as the number of stems, the plant volume or the emission potential differed significantly among the mono- or sesquiterpenoid classes. Hence, it is possible that morphological traits also play a role in mediating herbivore densities on individual plants. Previous studies suggested that both, the chemotype and the connected plant traits are crucial for host plant selection and performance of herbivorous insects. For instance, in *Brassica oleracea*, high levels of glucosinolates prolonged the development time of the specialist *Pieris rapae* and reduced survival in the generalist *Mamestra brassicae* (Gols et al., 2008). In *Salix sachalinensis*, leaf pubescence reduced overall leaf consumption by the willow leaf beetle *Melasoma lapponica* (Hayashi, Tahara & Ohgushi 2005). Furthermore, Carmona and colleagues (2010) showed that herbivore susceptibility depends on defence traits, including morphological and chemical traits. Our findings that sap-sucking aphids are influenced by terpenoid composition and height of the host plant, while ants are influenced by temperatures, support this general observation of the mediating role of plant chemical and morphological traits observed in other plant-insect systems, but also shows how the geographic location intermingles with these factors.

## Conclusion

We demonstrated that tansy chemotypes, defined by monoterpenoids or sesquiterpenoids, are not linked. Monoterpenoid chemotypes varied in a north-south direction, while sesquiterpenoid chemotypes were randomly clustered along the transect. In addition, we showed that monoterpenoid chemotype and the number of stems per plant influenced aphid occupancy, monoterpenoid chemotype affected ant occurrence. In contrast, aphid abundance was only influenced by the height of the host plant. Our findings validate and highlight the significance of the role of biogeography in shaping tansy chemical diversity with structuring effects for associated insect communities. Our results call for further analyses of the chemical signatures of plants in different habitat types to decipher how the environment selects for certain chemotypes and shapes their geographical distribution and how this selection consequently mediates plant-associated metacommunities.

## Supporting information

Supplement_Rahimova_et_al

## Acknowledgments

The authors thank Ina Zimmer, Kerstin Koch, and Baris Weber for technical support during the measurements. Additionally, the authors are grateful to fellow Ph.D. candidate Moritz Popp for his help on this work. This work was supported by grants of the Deutsche Forschungsgemeinschaft DFG within the Research Unit FOR3000 to JJR (JU2856/5-1), and WWW (WE3081/40-1) and joint DFG projects to (WWW (WE3081/25-2) and JPS (SCHN653/7-2).

## Contribution of authors

MC, MS, SZ, WWW, and JPS conceived and designed the study. MC, MS, and SZ did the sampling in Germany. MC performed the extraction, and YG conducted the GC-MS identification and quantification. HR and ANH analyzed the data with substantial input from HP, RRJ, LOP, RH, WWW, and JPS. HR and ANH wrote the initial version of the paper with contributions from HP, RH, and LOP. All authors read and approved the manuscript before submission.

## Data availability

The ecological and terpenoid data will be stored on DRYAD repository and made available after publication.

