## Supplement_Rahimova_et_al for "Geographic distribution of terpenoid chemotypes in *Tanacetum vulgare* mediates tansy aphid occurrence and abundance"

**ORCIDs:**

Humay Rahimova: 0009-0003-3453-6014

Annika Neuhaus-Harr: 0009-0009-6093-4108

Mary V. Clancy: 0000-0001-5597-4978

Yuan Guo: 0000-0002-5904-6113

Robert R. Junker: 0000-0002-7919-9678

Lina Ojeda-Prieto: 0009-0003-6069-901X

Hampus Petrén: 0000-0001-6490-4517

Matthias Senft: 0000-0003-3478-4774

Sharon E. Zytynska: 0000-0002-0174-3303

Wolfgang W. Weisser: 0000-0002-2757-8959

Robin Heinen: 0000-0001-9852-1020

Jörg-Peter Schnitzler: 0000-0002-9825-867X

**Supplementary figures:**

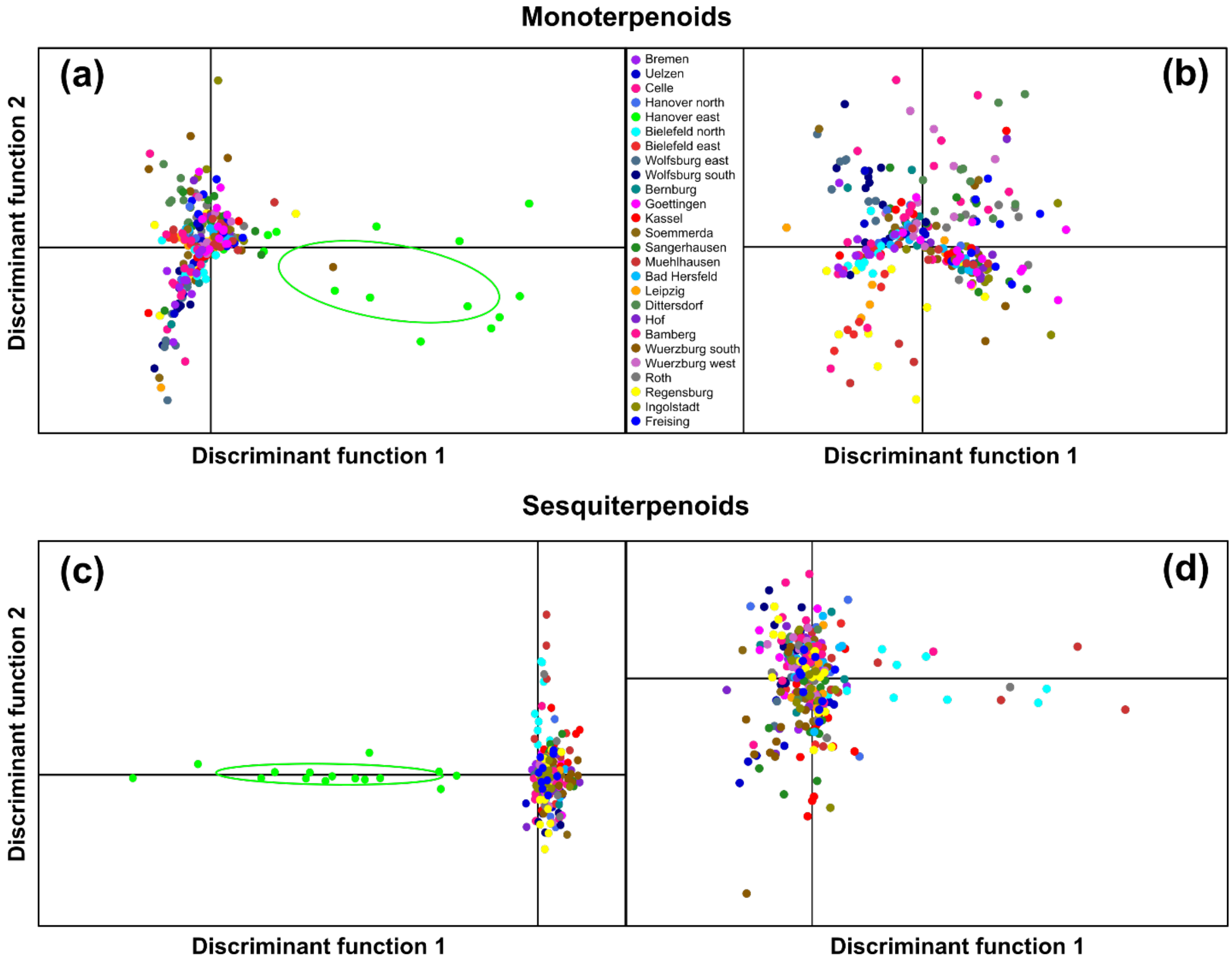


**Figure S1:** DAPC model of mono- and sesquiterpenoids with (a, c) and without (b, d) eastern Hanover site. The color of the dots resembles the collection site. Plants from the eastern Hanover site are highlighted by a green circle. When included they cluster outside of all other plants.

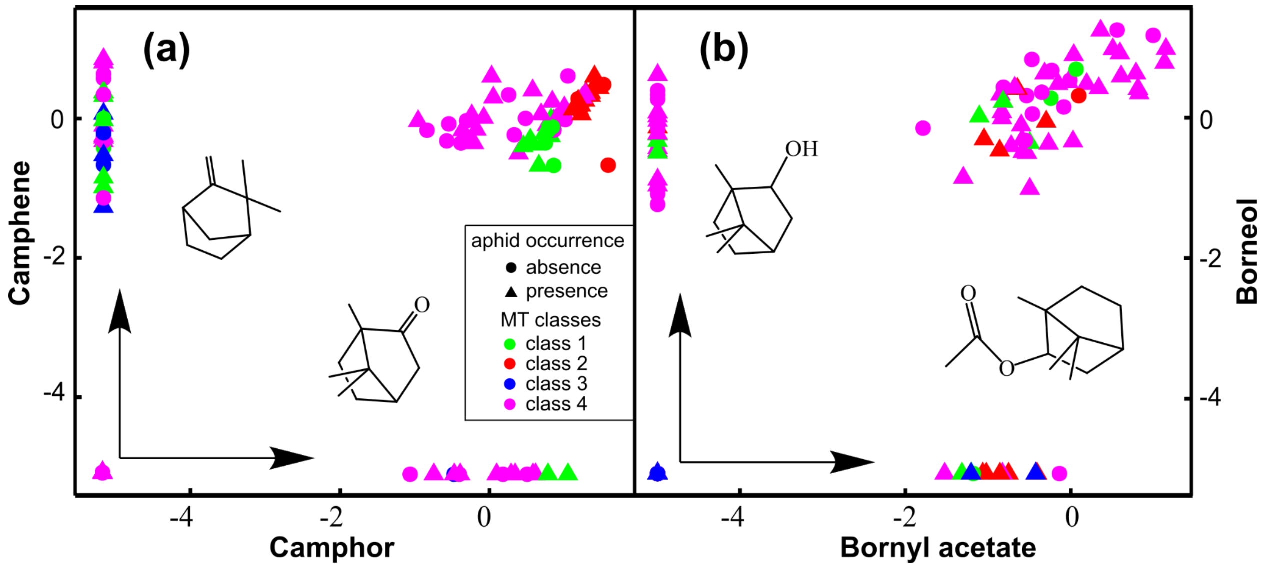


**Figure S2:** Association among the selected compounds in 278 plants. Panels show the relationship for the pairs of: (a) camphene and camphor, (b) borneol and bornyl acetate. Circles were used in plants without aphids, while triangles are used for plants with aphid presence.

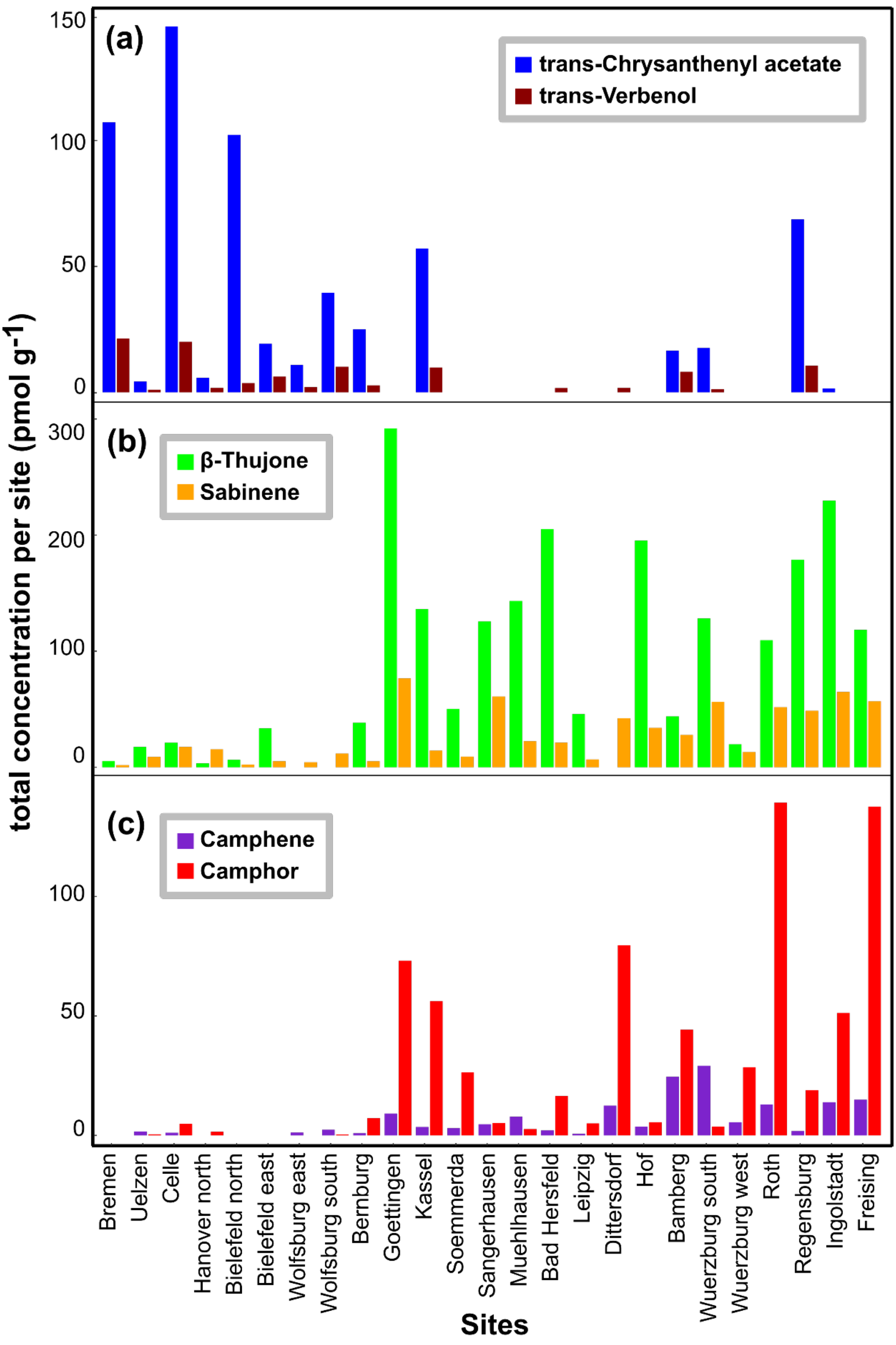


**Figure S3:** Distribution of the monoterpenoids across sampling sites from north to southern cities in Germany. Total concentration of (a) trans-chrysanthenyl acetate and trans-verbenol (b) β-thujone and sabinene (c) camphene and camphor per each site.

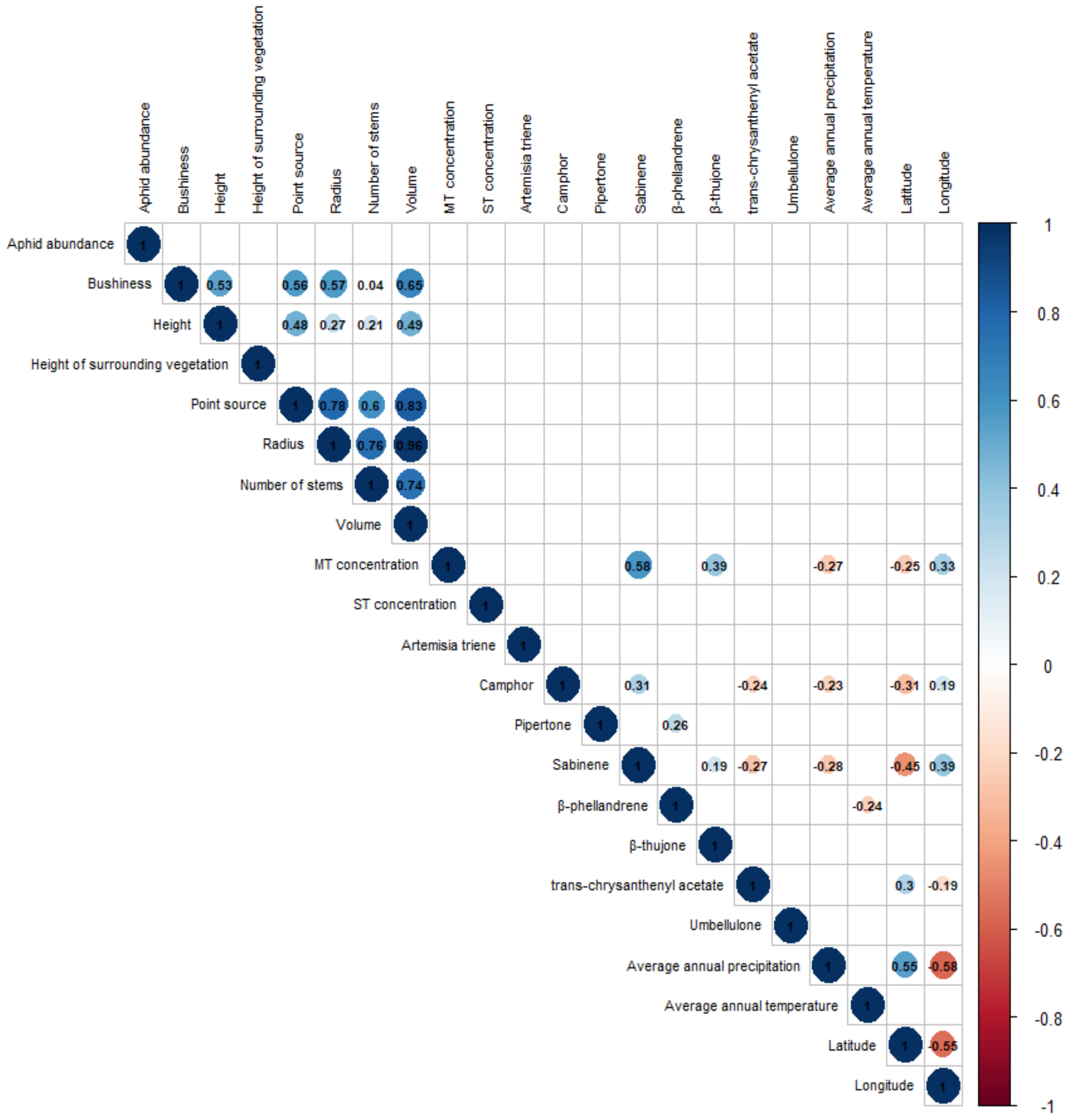


**Figure S4:** Correlation matrix of abundance of *M. fuscoviride*, all plant morphology variables, monoterpenoid (MT) and sesquiterpenoid (ST) concentrations, all monoterpenoid compounds exceeding a concentration of 25 pmol $g^{-1}$, average annual temperature (°C), average annual precipitation (mm), latitude, and longitude; Only significant values ($p < 0.05$) are shown (aphid and occurrence, and monoterpenoid and sesquiterpenoid classes were not included since these variables resemble factors and not values).

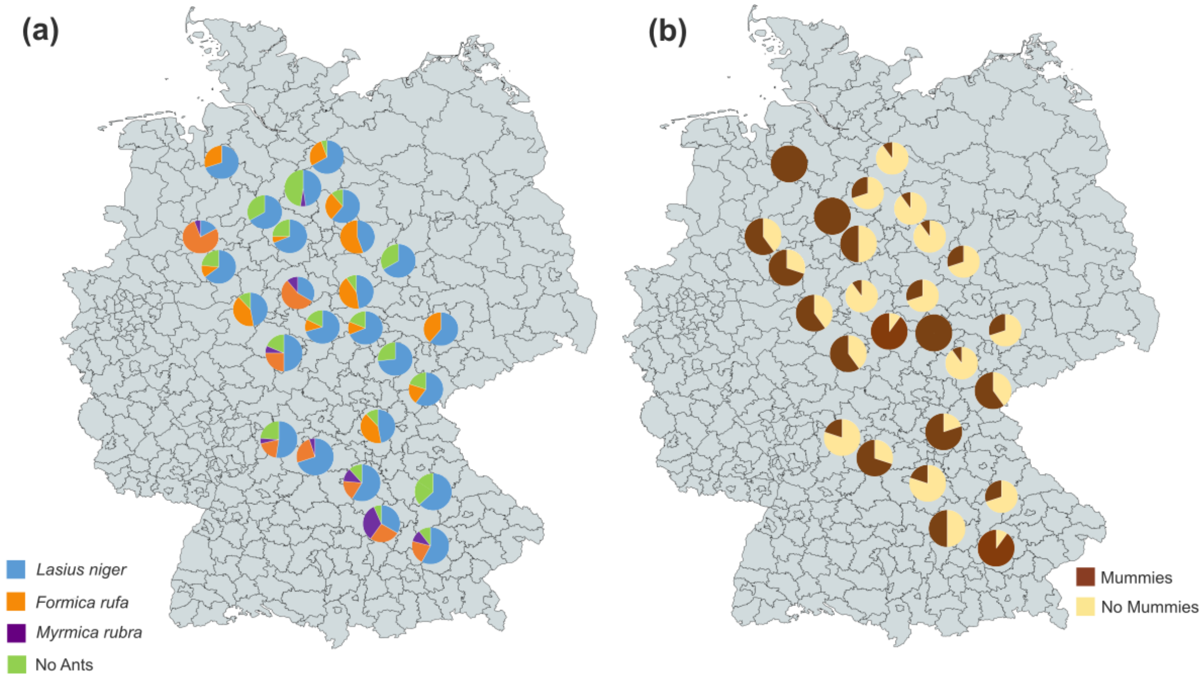


**Figure S5:** Presence of a) ant species and b) parasitized aphids across different sites across Germany. Note that more than one species could be found on the same tansy plant.

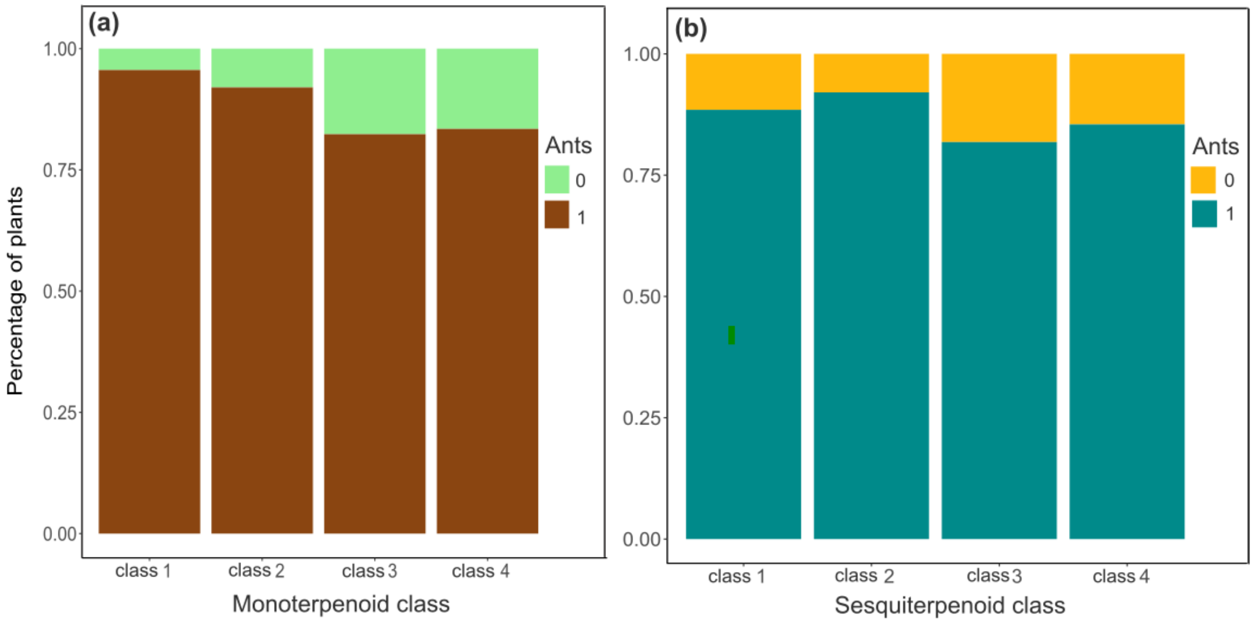


**Figure S6**: The bar charts indicate the percentage of numbers with ants (brown) and without ants (green) within the monoterpenoid classes (a) and with ants (blue) and without ants (orange) within in the sesquiterpenoid classes (b). A post-hoc Tukey test showed that ant occupancy in monoterpenoid class 1 was significantly higher compared to plants from monoterpenoid class 4 (std. error = 0.66, z-value = -2.65, p = 0.037).

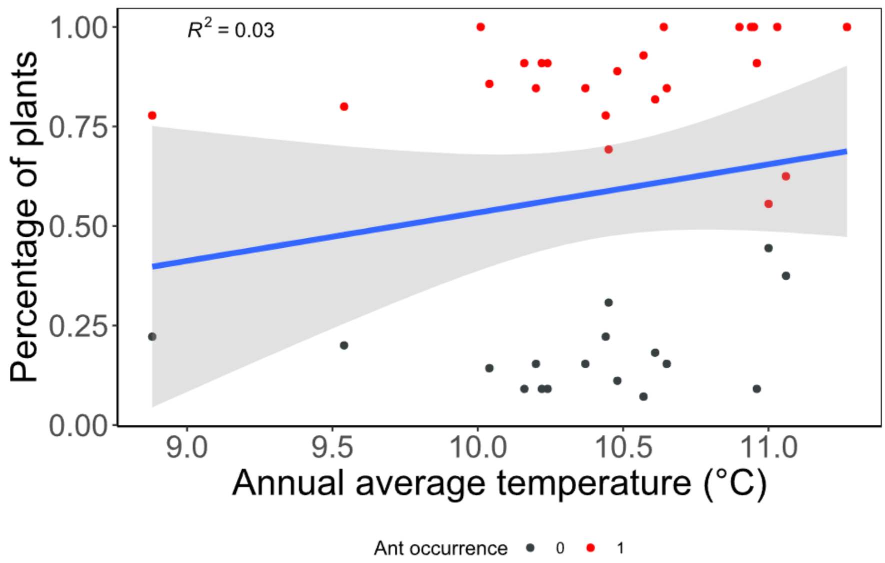


**Figure S7**: Presence and absence of ants across the different annual average temperatures. Ant presence is coded in red, ant absence in black.

**Supplementary tables:**

**Table S1:** Degrees of freedom (DF), statistic ($R^2$) and p-value for permutation test for adonis function (permutations = 999, method = "bray-curtis"). The test was conducted with the coordinate's variables (latitude and longitude) versus the monoterpenoid (MT) and sesquiterpenoid (ST) matrix data. Bold letters indicates significant p-values ($p < 0.050$) and italic letters indicates marginally significant values ($0.050 < p < 0.010$).

| **Coordinates** | **DF** | **MT matrix $R^2$ (p-value)** | **ST matrix $R^2$ (p-value)** |
|---|---|---|---|
| Latitude | 1 | **0.01 (<0.002)** | 0.002 (0.56) |
| Longitude | 1 | **0.01 (<0.001)** | 0.002 (0.58) |

**Table S2:** Degrees of freedom (DF), statistic (F-value) and p-value for plant trait variation across monoterpenoid and sesquiterpenoid classes using a one-factorial ANOVA. Bold letters indicates significant p-values (p <0.050) and italic letters indicates marginally significant values (0.05 < p <0.010).

| | DF | Monoterpenoid classes F (p-value) | Sesquiterpenoid classes F (p-value) |
|---|---|---|---|
| Number of stems | 3 | *2.55 (0.056)* | **3.69 (0.012)** |
| Plant volume | 3 | **2.71 (0.045)** | 1.73 (0.162) |
| Emission potential | 3 | **8.71 (<0.001)** | 0.74 (0.528) |
| Height | 3 | 1.45 (0.229) | 0.35 (0.792) |
| Radius | 3 | 2.00 (0.114) | 1.996 (0.120) |
| Bushiness | 3 | 1.02 (0.385) | 1.73 (0.162) |

**Table S3:** Degrees of freedom (DF), statistic (F-value) and p-value for plant traits variation across a latitudinal and longitudinal gradient in Germany using an one-factorial ANOVA. Bold letters indicates significant p-values (p < 0.050) and italic letters indicates marginally significant values (0.05 < p < 0.10).

| | | **Latitude** | **Longitude** |
|---|---|---|---|
| | **DF** | **F (p-value)** | **F (p-value)** |
| Number of stems | 1 | 1.35 (0.245) | 0.11 (0.74) |
| Plant volume | 1 | 0.38 (0.539) | 0.06 (0.808) |
| Emission potential | 1 | *3.19 (0.075)* | 1.38 (0.241) |
| Height | 1 | **10.20 (0.002)** | 0.04 (0.852) |
| Radius | 1 | 0.76 (0.383) | 1.86 (0.174) |
| Bushiness | 1 | 0.01 (0.931) | **8.23 (0.004)** |

**Table S4:** Degrees of freedom (DF), statistic ($X^2$ or F-value) and p-value for chemical, morphological, geographic, and biotic variables included, when applicable, in a binomial generalized linear model with aphid occurrence or ant occurrence as response variable, and in a GLM with aphid abundance as response variable. Bold letters indicates significant p-values ($p < 0.05$) and italic letters indicates marginally significant values ($0.05 < p < 0.01$).

| | **DF** | **Aphid presence $\chi^2$ (p-value)** | **Ant presence $\chi^2$ (p-value)** | **Aphid abundance F-value (p-value)** |
|---|---|---|---|---|
| MT class | 3 | **11.99 (0.007**)** | **10.62 (0.014*)** | 0.53 (0.661) |
| MT concentration | 1 | 0.56 (0.453) | 1.07 (0.301) | 0.44 (0.510) |
| ST class | 3 | 2.79 (0.426) | 2.16 (0.541) | 1.62 (0.186) |
| ST concentration | 1 | 0.31 (0.577) | 0.10 (0.756) | 1.48 (0.226) |
| MT class:ST class | 8 | 8.26 (0.409) | 10.08 (0.259) | 1.03 (0.414) |
| Emission potential | 1 | 0.38 (0.536) | 1.05 (0.306) | 0.31 (0.578) |
| Bushiness | 1 | 0.21 (0.643) | 2.21 (0.137) | 0.08 (0.783) |
| Height of surrounding vegetation | 1 | 0.10 (0.753) | 0.94 (0.332) | 2.58 (0.110) |
| Height | 1 | 1.85 (0.173) | 0.14 (0.706) | *3.17 (0.077)* |
| Stems | 1 | *3.76 (0.053)* | 0.63 (0.426) | 0.33 (0.568) |
| Mean annual temperature | 1 | 0.83 (0.362) | **3.91 (0.048*)** | 0.03 (0.854) |
| mean annual precipitation | 1 | 0.16 (0.694) | 0.61 (0.434) | 1.11 (0.293) |
| Latitude | 1 | 0.52 (0.471) | 0.53 (0.466) | 0.24 (0.624) |
| Longitude | 1 | 0.31 (0.577) | 0.01 (0.301) | 0.06 (0.805) |
| *Formica rufa* | 1 | - | - | 0.77 (0.381) |
| *Lasius niger* | 1 | - | - | 1.06 (0.304) |
| *Myrmica rubra* | 1 | - | - | 0.11 (0.745) |
| Residuals | 161 | - | - | |

**Table S5:** Output of binomial test for aphid presence in the different monoterpenoid and sesquiterpenoid classes. 95% confidence interval and p-values are reported. Bold letters indicates significant p-values ($p < 0.05$) and italic letters indicates marginally significant values ($0.05 < p < 0.01$).

| Aphid occurrence | Monoterpenoid classes | Sesquiterpenoid classes |
|---|---|---|
| | 95% confidence interval (p-value) | 95% confidence interval (p-value) |
| Class 1 | **0.52 - 0.82 (0.028)** | 0.22 - 0.69 (0.814) |
| Class 2 | 0.35 - 0.85 (0.454) | 0.41 - 0.73 (0.430) |
| Class 3 | 0.29 - 0.71 (1.000) | 0.18 - 0.90 (1.000) |
| Class 4 | 0.37 - 0.57 (0.617) | 0.44 - 0.62 (0.576) |

**Table S6:** Degrees of freedom (DF), statistic (F-value) and p-value for an analysis of variance (ANOVA) for *M. fuscoviride* occurrence and abundance compared to the functional Hill diversity (FHD) of monoterpenoids and sesquiterpenoids. Bold letters indicate significant p-values (p <0.05) and italic letters indicates marginally significant values (0.05 < p <0.01).

| | | **FHD MT** | **FHD ST** |
|---|---|---|---|
| | **DF** | **F (p-value)** | **F (p-value)** |
| Aphid occurrence | 1 | 1.49 (0.224) | 1.41 (0.236) |
| Aphid abundance | 1 | 1.29 (0.258) | 0.67 (0.412) |
